# Improved identification and differentiation from epileptiform activity of human hippocampal sharpwave-ripples during NREM sleep

**DOI:** 10.1101/702894

**Authors:** Xi Jiang, Jorge Gonzalez-Martinez, Sydney S. Cash, Patrick Chauvel, John Gale, Eric Halgren

## Abstract

In rodents, pyramidal cell firing patterns from waking may be replayed in NREM sleep during hippocampal sharpwave-ripples (HC-SWR). In humans, HC-SWR have only been recorded with electrodes implanted to localize epileptogenesis. Here, we characterize human HC-SWR with rigorous rejection of epileptiform activity, requiring multiple oscillations and coordinated sharpwaves. We demonstrated typical SWR in those rare HC recordings which lack interictal epileptiform spikes (IIS), and with no or minimal seizure involvement. These HC-SWR have a similar rate (~12/min) and apparent intra-HC topography (ripple maximum in putative stratum pyramidale, slow wave in radiatum) as rodents, though with lower frequency (~85Hz compared to ~140Hz in rodents). Similar SWR are found in HC with IIS, but no significant seizure involvement. These SWR were modulated by behavior, being largely absent (<2/min) except during NREM sleep in both stage 2 (~9/min) and stage 3 (~15/min), distinguishing them from IIS. This study quantifies the basic characteristics of a strictly selected sample of SWR recorded in relatively healthy human hippocampi.

## Introduction

Hippocampal (HC) sharpwave-ripples (SWR) are composed of a large negative ~50-100ms “sharpwave” with superimposed 80-200Hz ‘ripples’, followed by ~200ms positive delta-wave (Buzsáki, 2015). During SWR, hippocampal pyramidal cells “replay” spatio-temporal firing patterns from waking (Wilson and McNaughton, 1994). Memory is impaired by disruption of hippocampal replay (de Lavilléon et al., 2015), or of SWR (Ego-Stengel and Wilson, 2010; Suh et al., 2013), supporting SWR’s role in offline consolidation. The experiments above were in rodents, but SWR also occur in humans (Bragin et al., 1999a), in whom hippocampal lesions also disrupt memory consolidation (Squire et al., 2001). While there are differences between human and rodent SWR characteristics, there are also commonalities that could aide the identification and isolation of human SWR from electrophysiological recordings.

Rodent SWR occur in non-rapid eye movement sleep (NREM) and Quiet Waking periods of immobility with large amplitude EEG. Rodent sleep is mainly diurnal and highly fragmented (Van Twyver, 1969); human sleep is nocturnal and quasi-continuous, with distinct stages of N2 characterized by sleep spindles and K-complexes, and N3 characterized by rhythmic upstates and downstates (Silber et al., 2007). Rodents have a distinct behavioral state termed Quiet Waking with high amplitude waves (Roumis and Frank, 2015). Humans appear to lack this state. It is unknown if human SWR differ between N2 and N3, or if they occur during waking.

Although it is thus important to characterize SWR directly in humans, this requires HC recordings which can only be obtained in the context of clinical studies from patients implanted with electrodes localizing epileptogenic tissue. Consequently, literature values for basic characteristics of human SWR vary widely (Table 1). Human ripple center frequency ranges from ~85 Hz (Axmacher et al., 2008) to ~120 Hz (Bragin et al., 1999a), slower than rodents (~140 Hz), but similar to monkeys (Skaggs et al., 2007). However, clinical constraints in these studies hinder interpretation: some studies failed to separate interictal spikes from SWR (Clemens et al., 2007, 2011), or did not explicitly require that multiple oscillations be present in the ripple (Axmacher et al., 2008; Le Van Quyen et al., 2008; Nir et al., 2011; Brázdil et al., 2015), which could lead to single sharp transients being detected as well as sustained oscillations. Generally, studies did not require that a sharpwave be present, although in rodents and macaques many ripples occur without a sharpwave (Buzsáki, 2015; Ramirez-Villegas et al., 2015). Some studies recorded entirely (Clemens et al., 2007, 2011) or mainly (Axmacher et al., 2008; Le Van Quyen et al., 2008) outside the hippocampus, although high frequency oscillations in rodents vary considerably between regions (Buzsáki, 2015). No study distinguished anterior from posterior hippocampal recordings, although these differ considerably (Ranganath and Ritchey, 2012; Strange et al., 2014). Some studies recorded mainly or entirely outside of NREM stages N2 and N3 (Axmacher et al., 2008; Brázdil et al., 2015), perhaps because SWR occur in rodents during waking rest (Buzsáki, 2015), although humans may lack a similar state·.

**Table 1.** Previous studies reporting characteristics of human sharpwave-ripples.

| reference | location<sup>1</sup> | units | ripple selection |  |  |  |  |  | epilepsy rejection |  |  |  | ripple characteristics |  |  |  | #patients with ripples<sup>5</sup> | state<sup>6</sup> | similar references from same group |
| --- | --- | --- | --- | --- | --- | --- | --- | --- | --- | --- | --- | --- | --- | --- | --- | --- | --- | --- | --- |
|  |  |  | band pass | auto vs. visual<sup>2</sup> | minimum duration (ms) | min # waves | sharp wave required? | detection threshold<sup>3</sup> | exclude ictal onset zone? | exclude large sharp events? | exclude HC with IIS?<sup>4</sup> | visual control? | ripple frequency | ripple duration (ms) | sleep rate /min | wake rate /min |  |  |  |
| Axmacher et al, 2008 | HC, pHcG | no | 80-140 | a | 25 | 2 | no<sup>7</sup> | RLFP: 20μV | yes | yes | no | no | 85<sup>8</sup> | x | 0.35<sup>9</sup> | 5.3 | 11 | sleep, QW<sup>10</sup> |  |
| Bragin et al, 1999 | HC, pHcG | yes | 50-200 | v | 60 | x | no | SLFP: 100μV | no | no | no | yes | 120 | x | 5<sup>11</sup> |  | 4 | sleep, QW | Bragin et al, 1999 |
| Brázdil et al, 2015 | HC | no | 80-250 | av | 30 | 1 | no | RA: varied<sup>12</sup> | yes | no | no | no | x | 83 |  | 19.4 | 10 | waking task |  |
| Clemens et al, 2007 | pHCg (FO) | no | 80-140 | av | x | 3 | no | RA: varied<sup>12</sup> | no<sup>13</sup> | no | no | no | 84 | 27 | 1.3<sup>14</sup> | 0.85<sup>8</sup> | 10 | NREM (N2,N3) | Clemens et al, 2011 |
| Jacobs et al, 2016 | HC | no | 80-250 | v | x | x | no | unknown | yes<sup>15</sup> | yes | no | yes | x | x | 30<sup>16</sup> |  | 10 | NREM |  |
| Le Van Quyen et al, 2008 | HC, pHcG | yes | 80-200 | a | 20 | 1 | no | RA: 3 stdev | no | no | no | no | 98 | 38 | 20.7<sup>15</sup> |  | 11 | NREM, QW |  |
| Staba et al, 2004 | HC | no | 80-500 | a | no | 1 | no | RA: 3 stdev | yes<sup>15</sup> | no | no | no | 89 | x | 0.08<sup>8</sup> | 0.04 | 25 | NREM | Staba et al, 2002 |
| Staresina et al, 2015 | HC | no | 80-100 | a | 38 | 3 | no<sup>17</sup> | RMS: 99th pctl | yes | yes | no | yes | 87 | x | 1.2 |  | 12 | NREM (N2,N3) | Zhang et al, 2018 |
| This study | HC<sup>18</sup> | no | 60-120 | av | 25 | 4 | yes<sup>19</sup> | RMS: 80th pctl | yes | yes | yes<sup>20</sup> | yes | 82 | 96 | 12<sup>21</sup> | 1.9 | 20 | NREM (N2,N3) |  |

<sup>1</sup>HC-hippocampus; pHcG-parahippocampal gyrus (termed entorhinal or rhinal cortex by some); FO-recorded from cisterna ambiens with foramen ovale electrodes (all others are depth electrodes)
<sup>2</sup>av-automatic detection with visual validation; x-not reported
<sup>3</sup>RLFP-ripple LFP amplitude; SLFP-sharpwave LFP amplitude; RA-ripple bandpass-derived analytical signal amplitude; RMS: ripple bandpass-derived root mean square signal; stdev: number of standard deviations above the mean; pctl: percentile
<sup>4</sup>IIS-interictal spike
<sup>5</sup># of patients with ripples, in HC outside ictal onset zone, if reported separately
<sup>6</sup>QW-quiet waking
<sup>7</sup>ripple-triggered average shows small SW (~15μv)
<sup>8</sup>estimated from published figures
<sup>9</sup>N3 only
<sup>10</sup>almost all ripples were recorded in waking
<sup>11</sup>separate values not given for quiet waking
<sup>12</sup>scored by human reviewers
<sup>13</sup>ripples with vs without an IIS studied together
<sup>14</sup>average of N2 and N3

Here, we identified human SWR in HC without IIS, and use their characteristics to guide separation of SWR from IIS in other patients. Human SWR were similar to rodent SWR in waveform, rate, intra-hippocampal topography, and concentration in NREM sleep. They differ from rodents in their ripple frequency, with significantly different densities in N2/N3/waking and in anterior vs posterior HC. Overall, our more rigorous (than previous studies of human SWR) methods appear capable of isolating human SWR that are unlikely to be pathological in origin even from hippocampi with occasional IIS, given our SWR events’ similarities to rodent SWR and their distinct characteristics when compared to IIS events from the same patient population.

## Methods

### Patient selection

20 patients with long-standing drug-resistant partial seizures underwent SEEG depth electrode implantation in order to localize seizure onset and thus direct surgical treatment (see Table 2 for demographic and clinical information). Patients were selected from a group of 54 based on the following criteria: (1) age between 16 and 60 years; (2) globally typical SEEG rhythms in most channels (i.e., absence of diffuse slowing, widespread interictal discharges, highly frequent seizures, etc.); (3) electrode contacts in the HC as verified using non-invasive imaging (see below); (4) no previous excision of brain tissue or other gross pathology; (5) at least one HC contact in an HC not involved in the initiation of seizures. The resulting group of 20 patients includes 6 patients with an HC contact in a location with no interictal spikes, which were used to guide the protocols applied to the remaining 14 patients (Table 2). The 20 patients included 7 males, aged 29.8±11.9 years old (range 16-58). Electrode targets and implantation durations were chosen entirely on clinical grounds (Gonzalez-Martinez et al., 2013). All patients gave fully informed consent for data usage as monitored by the local Institutional Review Board, in accordance with clinical guidelines and regulations at Cleveland Clinic.

**Table 2.** Patient characteristics. Pt: patient. L: Left. R: Right. Dur.: duration. Std dev: standard deviation.

| Pt# | Age | Sex | Handedness | L aHC | L pHC | R aHC | R pHC | # Cortical Channels | Interictal-free? | # Sleep Periods | mean duration of | Std dev NREM (h) | Total N2 Dur. (h) | Total N3 Dur. (h) | SWR Rate, N2 aHC (/min) | SWR Rate, N2 pHC (/min) | SWR Rate, N3 aHC (/min) | SWR Rate, N3 pHC (/min) |
| --- | --- | --- | --- | --- | --- | --- | --- | --- | --- | --- | --- | --- | --- | --- | --- | --- | --- | --- |
| 1 | 20 | M | R | x |  |  |  | 19 | N | 3 | 2.2 | 0.78 | 2.8 | 3.3 | 11 |  | 19 |  |
| 2 | 51 | F | R | x |  |  |  | 12 | N | 4 | 7.5 | 1.87 | 27.3 | 0 | 1.6 |  |  |  |
| 3 | 58 | F | R | x | x |  |  | 24 | N | 4 | 7.1 | 2.44 | 23.1 | 0.30 | 11 | 12 | 24 | 25 |
| 4 | 42 | M | L | x | x | x |  | 17 | N | 4 | 3.1 | 0.33 | 1.8 | 9.0 | 10 | 4.3 | 11 | 4.3 |
| 5 | 18 | F | L |  |  | x | x | 21 | Y | 1 | 3.7 |  | 1.9 | 0.9 | 21 | 3.7 | 38 | 14 |
| 6 | 20 | F | R |  |  | x | x | 21 | N | 3 | 2.7 | 1.49 | 2.6 | 3.0 | 9.1 | 12 | 15 | 17 |
| 7 | 22 | M | LR | x |  |  |  | 19 | Y | 3 | 3.8 | 1.17 | 2.4 | 3.8 | 11 |  | 13 |  |
| 8 | 30 | F | R |  |  | x |  | 13 | N | 5 | 5.2 | 1.26 | 6.5 | 14 | 19 |  | 20 |  |
| 9 | 43 | F | R | x | x |  |  | 12 | N | 4 | 3.1 | 0.45 | 3.7 | 4.4 | 13 | 7.9 | 16 | 7.2 |
| 10 | 16 | M | R |  |  | x |  | 17 | Y | 5 | 3.8 | 1.11 | 7.5 | 8.8 | 13 |  | 18 |  |
| 11 | 32 | F | R | x |  | x | x | 30 | N | 3 | 5.1 | 3.43 | 8.4 | 2.8 | 7.3 | 7.7 | 12 | 12 |
| 12 | 36 | M | L |  |  |  | x | 25 | Y | 4 | 5.2 | 1.48 | 11.6 | 6.5 |  | 5.5 |  | 7 |
| 13 | 21 | F | L | x | x | x |  | 14 | N | 3 | 6 | 0.47 | 14.7 | 1.1 | 13 | 6.2 | 19 | 4.9 |
| 14 | 21 | F | R |  | x |  |  | 14 | N | 8 | 3.7 | 1.1 | 16.9 | 9.3 |  | 11 |  | 15 |
| 15 | 29 | F | R |  | x |  | x | 17 | N | 4 | 2.5 | 1.31 | 5.4 | 2.7 |  | 3.6 |  | 7.2 |
| 16 | 41 | F | R |  |  | x |  | 19 | N | 3 | 4.6 | 0.72 | 7.4 | 4.4 | 9.4 |  | 11 |  |
| 17 | 24 | M | R | x |  |  |  | 21 | N | 3 | 5.1 | 1.88 | 8.9 | 2.9 | 23 |  | 24 |  |
| 18 | 31 | F | R | x |  |  |  | 15 | N | 6 | 6.1 | 1.18 | 24.2 | 3.8 | 7.8 |  | 10 |  |
| 19 | 21 | M | R | x | x |  |  | 15 | Y | 4 | 2.8 | 0.72 | 5.4 | 5.8 | 12 | 2.6 | 19 | 6.1 |
| 20 | 19 | F | R |  |  | x |  | 21 | Y | 3 | 4.9 | 2.53 | 5.2 | 6.0 | 11 |  | 12 |  |
| **Mean** | **30** |  |  |  |  |  |  | **18** | **Total: 8** | **4** | **4.4** |  | **9.39** | **4.64** | **12** | **7** | **18** | **11** |

### Electrode localization

After implantation, electrodes were located by aligning post-implant CT to preoperative 3D T1-weighted structural MRI with ~1mm^3^ voxel size (Dykstra et al., 2012), using 3D Slicer (RRID:SCR_005619). This allows visualization of individual contacts with respect to HC cross-sectional anatomy (e.g., Fig. 1*F*), which was interpreted in reference to the atlas of Duvernoy (Duvernoy, 1988). At most anterior-posterior levels, an electrode approaching human HC orthogonal to the sagittal plane passes through white matter, ventricle, alveus, and strata oriens, pyramidale, radiatum and lacunosum-moleculare, in that order. This progression was correlated anatomically by superposition of electrode contacts from post-implant CT onto preoperative MRI, and physiologically by observing activity typical of white matter, CSF or gray matter.

**Figure 1.**
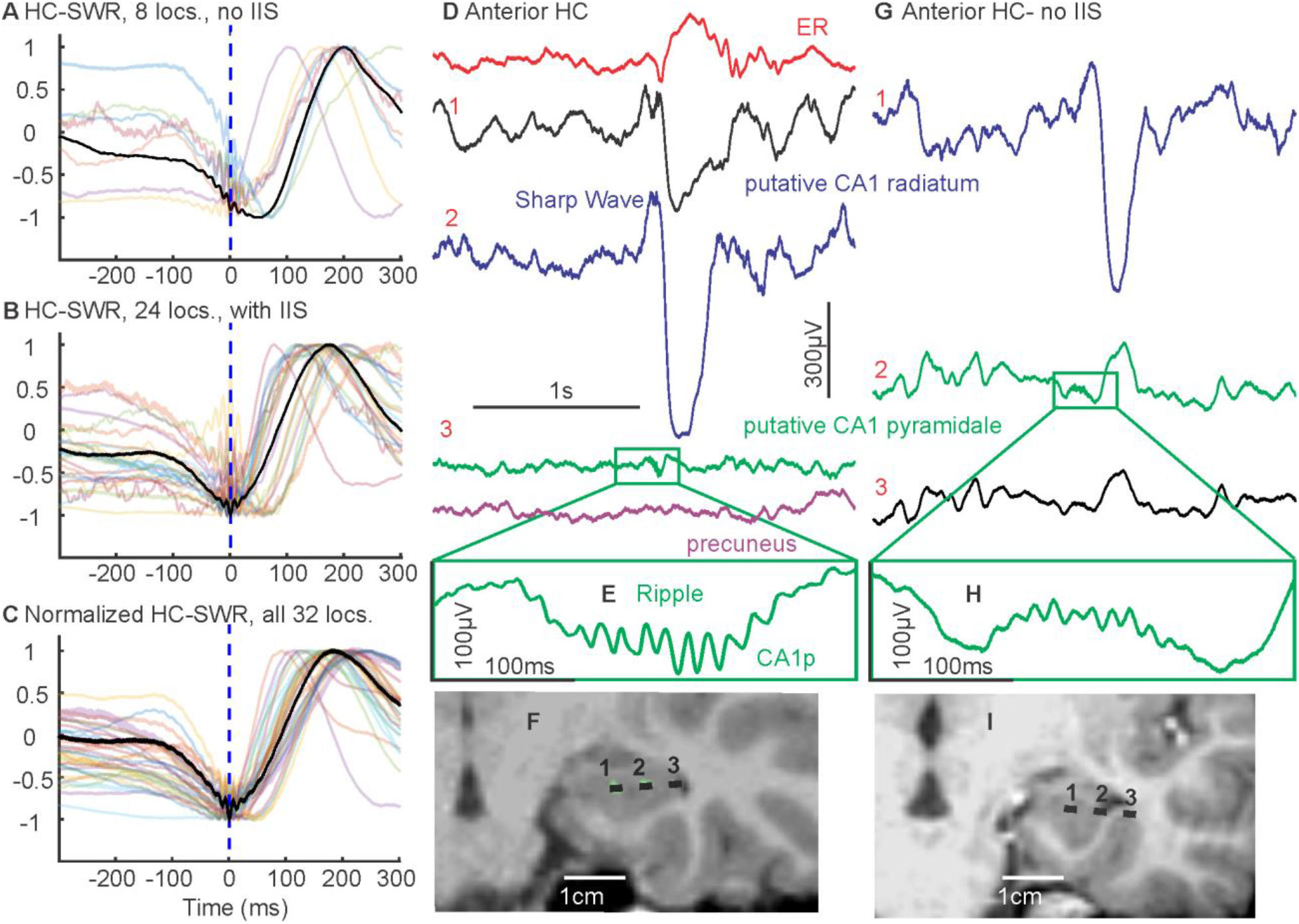
Distribution and LFP characteristics of HC-SWR. ***A-C***. Overlaid average waveforms of SWR across hippocampal contacts from different patients, with one representative patient bolded in black for clarity. ***A***. Hand-marked SWR exemplars from interictal-free patients. ***B***. Hand-marked SWR exemplars from other patients. ***C***. Automatically detected SWR from all patients. ***D***. Single sweep of HC and NC referential SEEG showing SWR. ***E***. Expanded time-base of ripple. ***F***. MRI with CT of SEEG contacts in ***D*** superimposed. ER: entorhinal. ***G***-***I*** are replicates of ***D***-***F***, but from another subject with no HC interictal activity. locs.: hippocampal recording sites.

The assignment of depth contacts to anterior or posterior hippocampus (aHC/pHC) was made with the posterior limit of the uncal head as boundary (Poppenk et al., 2013; Ding and Van Hoesen, 2015). Recordings were obtained from 32 HC contacts, 20 anterior (11 left) and 12 posterior (7 left). In 4 patients, HC recordings were bilateral (3 anterior and 1 posterior), and in 8 patients, ipsilateral anterior and posterior HC were both recorded. The distance of each hippocampal contact from the anterior limit of the hippocampal head (e.g., Fig. 3*F*) was obtained in Freesurfer (RRID:SCR_001847).

### Data collection and preprocessing

Continuous recordings from SEEG depth electrodes were made with a cable telemetry system (JE-120 amplifier with 128 or 256 channels, 0.016-3000 Hz bandpass, Neurofax EEG-1200, Nihon Kohden) across multiple nights (Table 2) over the course of clinical monitoring for spontaneous seizures, with 1000 Hz sampling rate. The total NREM sleep durations vary across patients; while some differences are expected given the intrinsic variability of normal human sleep duration (Carskadon and Dement, 2010), as well as the sleep disruption inherent in the hospital environment, we identified a subset of 28 sleep sessions across 16 of our patients whose N2 and N3 durations were comparable to normative data (i.e., within 2 standard deviations) (Moraes et al., 2014). Supplementary analyses of these sleep sessions were used to confirm that they exhibited the basic results seen in the entire group.

Recordings were anonymized and converted into the European Data Format (EDF). Subsequent data preprocessing was performed in MATLAB (RRID:SCR_001622); the Fieldtrip toolbox (Oostenveld et al., 2011) was used for bandpass filters, line noise removal, and visual inspection. Separation of patient NREM sleep/wake states from intracranial LFP alone was achieved by previously described methods utilizing clustering of first principal components of delta-to-spindle and delta-to-gamma power ratios across multiple LFP-derived signal vectors (Gervasoni et al., 2004; Jiang et al., 2017), with the addition that separation of N2 and N3 was empirically determined by the proportion of down-states that are also part of slow oscillations (at least 50% for N3 (Silber et al., 2007)), since isolated down-states in the form of K-complexes are predominantly found in stage 2 sleep (Cash et al., 2009).

### HC-SWR identification (Fig. 2H)

Previous studies have used a variety of methods to identify SWR and distinguish them from IIS (Table 1). Previous studies did not require that a sharpwave be present. However, based on the fact that in rodents and macaques many ripples occur without a sharpwave (Buzsáki, 2015; Ramirez-Villegas et al., 2015), we explicitly tested for the presence of a sharpwave and following slow positivity, with the ripple needing to occur near the peak of the sharpwave. In order to identify ripples, previous studies detected outlier peaks in the amplitude or power in the putative ripple frequency range. However, such peaks can reflect a single sharp transient. Thus, unlike some previous studies (Table 1), in our study we followed detections of ripple-frequency amplitude peaks with tests for multiple discrete peaks at ripple frequency. Since IIS are characterized by a large fast spike component, the test for multiple oscillations helps distinguish them from SWR. Since the slower field potentials accompanying IIS have different waveforms than the characteristic sharpwave and later positivity, requiring that these waves be present in their typical form also helps differential IIS from SWR. Nonetheless, due to the variability of IIS, it is important to visually examine the recordings in each patient to confirm the adequacy of IIS-rejection. These steps have seldom been taken by previous studies, some of which failed to even attempt to separate interictal spikes from SWR (Clemens et al., 2007, 2011). Some studies recorded entirely (Clemens et al., 2007, 2011) or mainly (Axmacher et al., 2008; Le Van Quyen et al., 2008) outside the hippocampus, although high frequency oscillations in rodents vary considerably between regions (Buzsáki, 2015). In our study, we recorded SWR exclusively from contacts which were confirmed to lie in the HC with CT/MRI localization. This localization was also used in our study to distinguish anterior from posterior HC recordings, which has not previously been done. This distinction is important given that these areas differ considerably in functional correlates and external anatomical connections (Ranganath and Ritchey, 2012; Strange et al., 2014). Finally, although most studies recorded human SWR in NREM sleep stages N2 and N3, some studies recorded mainly or entirely in other states, especially waking (Axmacher et al., 2008; Brázdil et al., 2015). Since it is unclear whether these are the same phenomena as classical SWR, we recorded during repeated 24-hour periods and after determining that SWR are essentially absent outside to N2 and N3, confined our quantitative analysis to those sleep stages.

In summary, we attempted to identify HC-SWR which were distinct from IIS and which possessed the ripple, sharpwave, and positive wave components of the SWR (Fig. 2*H*). Initially, an inclusive selection criterion (ripple selection based on amplitude from 65-180 Hz) was applied to HC recordings in the patients without IIS described above, in order to establish the parameters appropriate to use in the complete patient population. A minimum of 300 SWR were selected visually based on waveform and averaged in the time and time-frequency domains. Based on the ripple frequency observed in this population, a bandpass of 60 to120 Hz was chosen for further processing (6th order Butterworth IIR bandpass filter, with zero-phase forward and reverse filtering). Root-mean-square (RMS) over time of the filtered signal was calculated using a moving average of 20 ms, with the 80th percentile of RMS values for each HC channel being set as a heuristic cut-off. Whenever a channel’s signal exceeds this cut-off, a putative ripple event was detected. Adjacent putative ripple event indices less than 40ms apart were merged, with the center of the new event chosen by the highest RMS value within the merged time window. Each putative ripple event was then evaluated based on the number of distinct peaks in the HC LFP signal (low-passed at 120 Hz) surrounding the event center; a 40-ms time bin was shifted (5ms per shift) across ±50 ms, and at least one such time bin must include more than 3 peaks (the first and the last peak cannot be both less than 7ms away from the edges of this time bin) for the putative ripple event to be considered for subsequent analyses. In addition, the distance between two consecutive ripple centers must exceed 40 ms. To determine the duration of each ripple, the left edge was defined as the first time point before the ripple center where RMS value exceeded the percentile threshold used for initial ripple detection, and the right edge was defined as the first time point after the ripple center where RMS value fell below the threshold; all other time points within the duration of this ripple must exceed the RMS threshold.

To identify which ripples were coupled to sharpwaves (i.e., SWR), we created patient-specific average templates (−100ms to +300ms around ripple center) from hand-marked exemplars (100-400 SWR) in NREM per patient) that resemble previously described primate sharpwave-ripples with biphasic LFP deflections (Skaggs et al., 2007; Ramirez-Villegas et al., 2015). We then evaluated whether each ripple qualifies as a SWR with the following two criteria: 1, the similarity of the peri-ripple LFP to the average template as quantified by the dot product between the template and the peri-ripple LFP (−100ms to +300ms). For each template-LFP pair, the similarity threshold was chosen so that it would reject at least 95% of the hand-marked ripples with no sharpwave. 2, the absolute difference between the LFP value at the ripple center and at the maximum/minimum value between +100ms and +250ms after ripple center was computed for each ripple; this difference must exceed 10% (or another value which excludes 95% of hand-marked ripples with no sharpwave) of the difference distribution created from hand-marked SWR. In addition, the distance between two consecutive SWR centers must exceed 200 ms. Time-frequency plots of HC LFP centered on detected SWR were then created in MATLAB with EEGLAB toolbox (Delorme and Makeig, 2004), each trial covering ±1500 ms around SWR and with -2000 ms to -1500ms as baseline (masked for significance at alpha = 0.05, 200 permutations).

Since RMS peaks may arise from artifacts or epileptiform activity, 2000 ms of hippocampal LFP data centered on each ripple event undergoes 1-D wavelet (Haar and Daubechies 1-5) decomposition for the detection and removal of sharp transients (i.e. signal discontinuities). For each wavelet decomposition, putative SWR were marked in ~10 min long NREM sleep period (marking ended sooner if 400 putative SWR had been found). A scale threshold was then established via iterative optimization for the best separation between hand-marked true ripple events and interictal events in the same NREM sleep period. Each putative sharp transient was then rejected only if the 200 Hz highpassed data at that point exceeds an adaptive threshold (Bragin et al., 1999a) of 3—or another number that allows best separation in agreement with visual inspection by human expert (between 0.5 and 5, and ~2 on average)—standard deviations above the mean for the 10-second data preceding the transient.

We also compared the occurrence rates and waveform peak-to-peak LFP amplitudes of SWR in patients without IIS to those in other patients, using linear mixed effect models (implemented in MATLAB as part of the Statistics and Machine Learning Toolbox). Specifically, for each HC-SWR in NREM, we created three separate models per response variable (i.e. SWR occurrence rates or waveform amplitudes): a baseline model where responses are predicted by whether a given hippocampal site is IIS-free (Response ~ 1 + Patient Category), a model including patient random effect as a random intercept term (Response ~ 1 + Patient Category + (1 | Patient ID)), and a model with patient random effect as both random intercept and slope terms (Response ~ 1 + Patient Category + (Patient Category | Patient ID)). Comparisons between model fits were conducted with likelihood ratio tests, with the random intercept models being the best fit for rate-response, and the baseline model being the best fit for amplitude-response.

### Code Accessibility

All custom scripts would be available upon request by contacting the corresponding author and would be delivered through the UCSD RDL-share service.

## Results

We identified human HC-SWR from intracranial recordings, and distinguished them from IIS by characterizing their density, topography, and spectral traits. Morphologically normal ripples were isolated from >24 h continuous recordings in 20 stereoelectroencephalography (SEEG) patients with intractable focal epilepsy, with anterior (in 17 patients) and/or posterior HC contacts (in 11 patients).

### Comparison of SWR in patients with or without HC IIS

Given the potential for contamination with epileptiform activity, our initial studies focused on six patients (8 HC locations, 5 in aHC, 3 in pHC) that show no HC interictal spikes (IIS) over the course of the patient’s hospitalization (Table 2). SWR were detected by applying patient-specific SWR templates built from hand-marked represenattives to non-epileptiform high frequency oscillation events, we found that these patients showed SWR with the same characteristic waveform, rate (for anterior contacts, ~13.45/min in N2, ~19.89/min in N3, comparable to some previous reports, see Table 1), and association with NREM as reported in rodents and in non-human primates (Fig.1*A*, Fig. 2*A*, Fig. 3*A*). These characteristics were compared to those obtained in patients whose HC contacts showed occasional IIS. Events in these patients were accepted as SWR only after rigorous separation procedures from IIS based on amplitude, waveform, and spectral pattern (Fig. 2*A*, *B*, *C*, *G*). The SWR in the patients with HC IIS were comparable to those with no IIS in waveform and rate (Fig. 1*A*-*C*, Fig. 3*F*-*G*). Further studies then included these patients and HC electrodes (24 unique contacts, 32 total including IIS-free patients). We also compared the occurrence rates and peak-to-peak LFP amplitudes of HC-SWR in IIS-free patients with those in other patients, using linear mixed effect models (with patient ID being a random effect) to evaluate the significance of IIS contamination of HC on SWR characteristics in NREM. We found no significant effect of patient category (i.e. IIS-free or not) regarding either SWR rates or amplitudes (p > 0.7852 and p > 0.1712, respectively).

**Figure 2.**
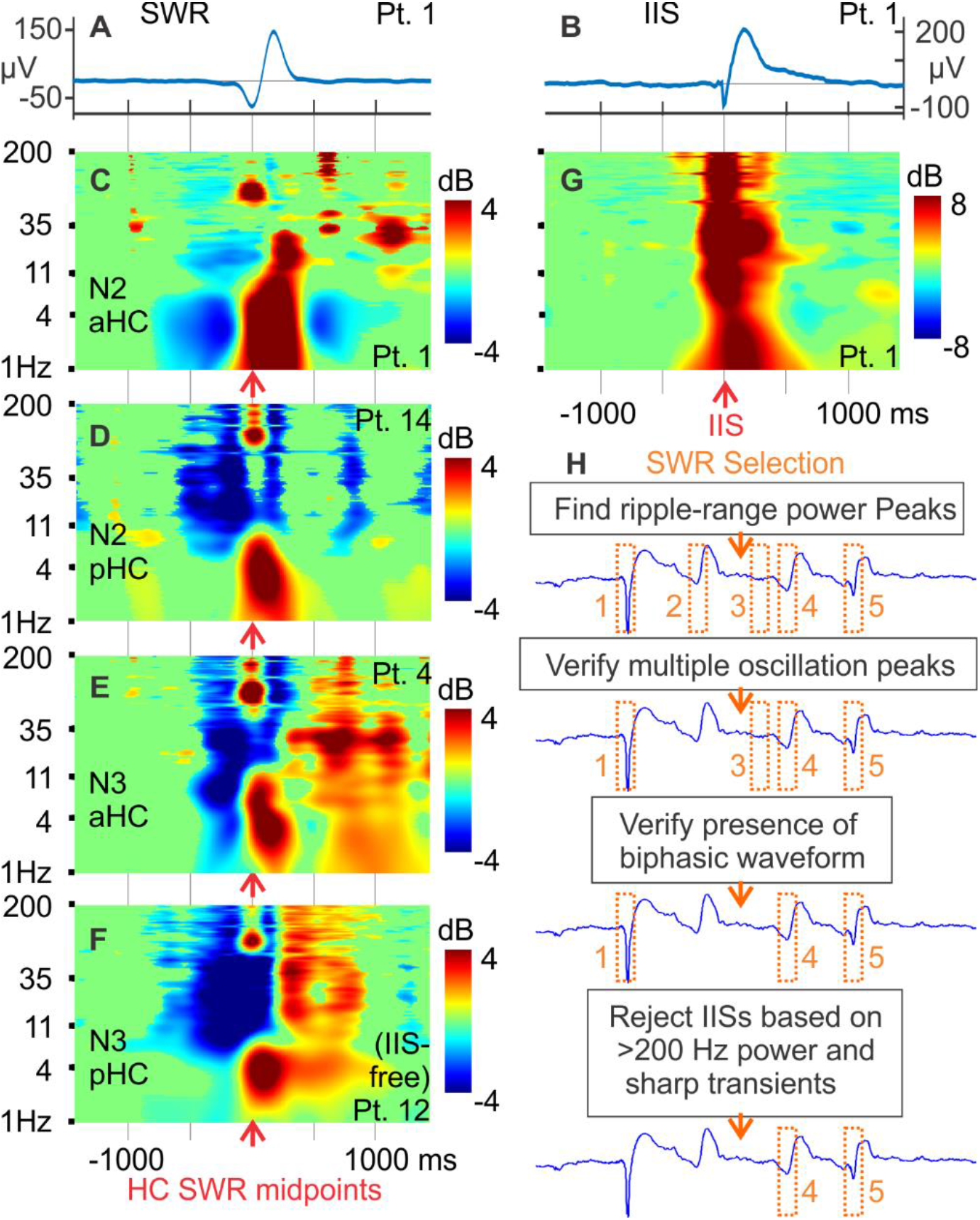
Spectral characteristics of HC SWR. ***A***. Average LFP of SWR accepted after the procedure in ***H***. ***B***. Average LFP of IISs originally classified as SWR based on high-frequency power from the same HC contact as ***A***, but rejected from further analysis. ***C-F***. Time-frequency plots showing typical frequency ranges of SWR, and broadband power decreases near SWR, from different NREM stages and example contacts in different HC locations. ***G***. time-frequency plot of the same IISs that make up ***B***; the power increase across all frequencies is not characteristic of true SWR (see ***C***, which was derived from true SWR in the same HC contact). ***H***. Schematic for the identification process of true SWR. Candidate LFP events (examples marked 1-5 and bound in orange dash line boxes) were first selected based on 60-120 Hz power. Putative ripples resembling event 2, i.e. those with less than 3 unique oscillation peaks, would be rejected first, followed by the removal of instances resembling event 3 (no distinct sharpwave that resembles hand-marked examples) and of those similar to event 1 (putative interictal spikes with high frequency power above ripple range).

**Figure 3.**
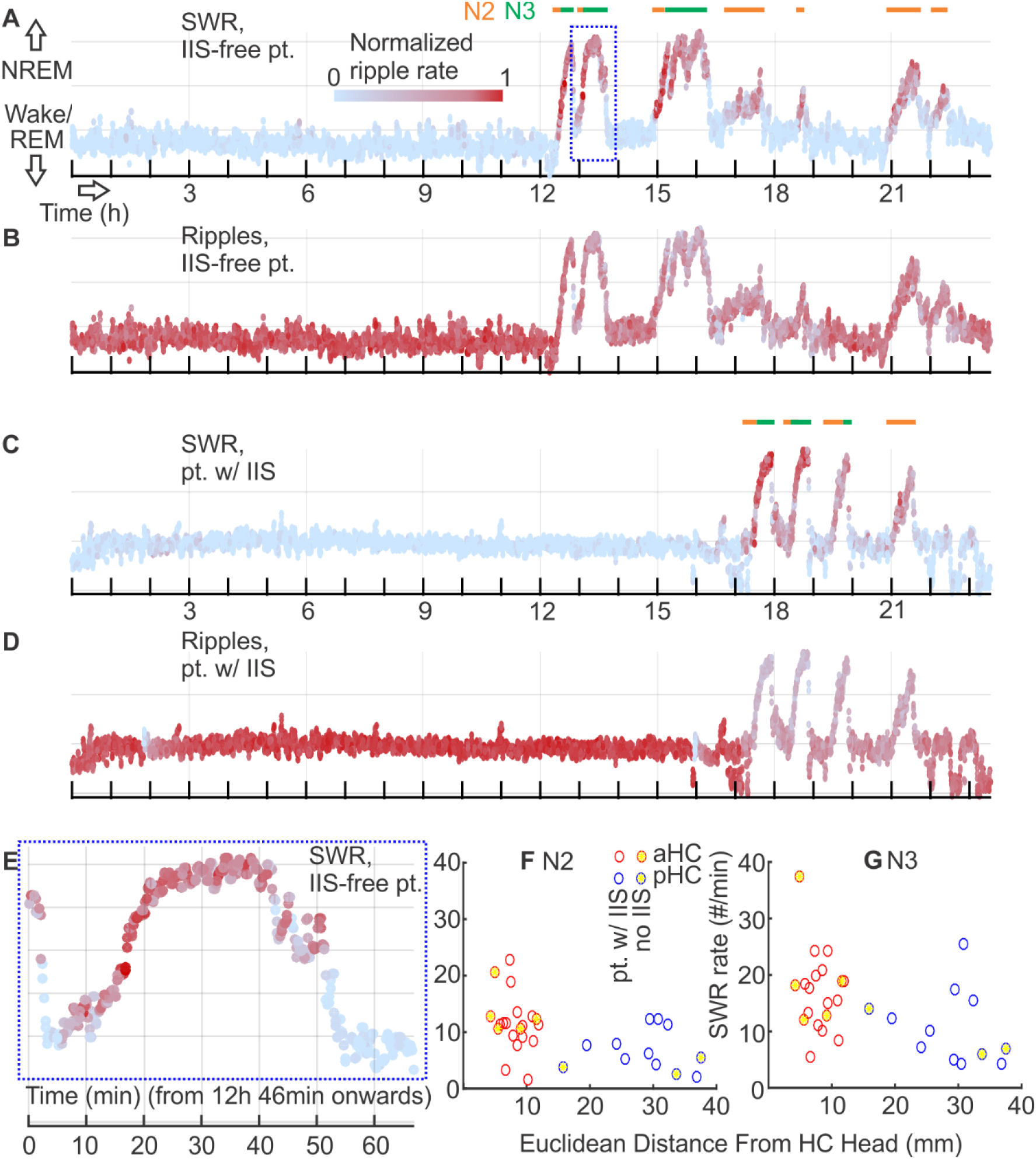
Distribution of HC SWR across stages of sleep and waking. ***A-D***. State plots showing the separation of NREM sleep in 24-hour LFP recording for both IIS-free (***A***, ***B***) and non-IIS-free (***C***, ***D***) patients, using the first principal component derived from 19 vectors (one per cortical bipolar SEEG channel) of frequency power ratios (0.5-3 Hz over 0.5-16 Hz) (Gervasoni et al., 2004; Jiang et al., 2017). SWR rate (***<u>A</u>***, ***C***) and plain ripple rate (***B***, ***D***) are color coded with red intensity (normalized within each subject), and N2/N3 periods are marked with orange/green horizontal lines, respectively. Deep blue dash line in ***A*** marks the close-up period in ***E***.***E***, close-up of one NREM cycle in ***A***. Each data point in ***A***-***E*** covers 30 seconds, with 10-second overlap between two adjacent points. ***F***, ***G***. Distribution of SWR occurrence rates in different NREM stages across the HC longitudinal axis. Each circle represents data from one hippocampal contact. Red circles indicate contacts in aHC; blue circles indicate those in pHC. Yellow fill marks the contacts from interictal-free patients.

### Intrahippocampal SWR topography and ripple frequency

We observed in our data that the intrahippocampal topography of the SWR in humans appears similar to that established in rodents, for both IIS-free and occasional-IIS HC: the maximum amplitude of the ripple is seen just after passing through the alveus into the presumptive stratum pyramidale, and that of the sharpwave and following delta wave was maximal more medially in presumed stratum radiatum (Fig. 1*D*-*I*). We further quantified the relative maximum amplitude of all ripples (that were part of SWR) and of their associated sharp-delta waves in radiatum versus pyramidale contacts in 10 subjects with contacts that putatively sampled both layers. For ripples, we computed for each SWR the proportion of increase for maximum LFP amplitude over mean LFP amplitude across ±20ms of ripple center; for sharp-delta wave, we computed for each SWR the proportion of increase for maximum LFP amplitude over mean LFP amplitude across 1000 ms (−250~750 ms relative to ripple center). We found that the mean pyramidale relative ripple amplitude (0.68) was significantly greater than the mean radiatum relative ripple amplitude (0.63) (p < 2.2 × 10^−16^, paired two-tailed t-test), although the effect size was small (Cohen’s *d* = 0.05) (Lakens, 2013). As expected from examples in Fig. 1*D* and Fig. 1*G*, the mean radiatum relative sharp-delta wave amplitude (3.7) was significantly greater than the mean pyramidale relative sharp-delta wave amplitude (2.2) (p < 2.2 × 10^−16^, paired two-tailed t-test), with a medium effect size (Cohen’s *d* = 0.54).

Time-frequency plots of SWR showed a concentration of ripple power at the oscillation frequency (Fig. 2*C*-*F*). The mean frequency of ripple oscillations in detected SWR was 81.5±9.7 Hz. This is lower than observed in rodents, but is consistent with previous studies in primates (Skaggs et al., 2007; Logothetis et al., 2012).

### SWR density in different stages of NREM sleep and in waking

Consistent with the occurrence of SWR during slow wave sleep in rodents, human SWR were largely confined to NREM sleep (Fig. 3*A*, *C*). The mean occurrence rate of SWR was significantly greater in anterior hippocampus (aHC) than in posterior hippocampus (pHC), and more SWR occur in N3 than in N2 (Fig. 3*F*-*G*, Table 2). In contrast to NREM, SWR density was low in waking (Fig. 3*A*, *C*) for aHC (1.77/min) and pHC (2.16/min), which did not differ significantly (p = 0.5433, two-tailed two-sample t-test).

Patients in the hospital environment typically undergo varying degrees of sleep disruption. In order to evaluate if our results were influenced by this disruption, we compared basic SWR parameters in patients with relatively normal sleep to our total group. N2 and N3 durations both fell within 2 standard deviations of normative data across 12 of our patients (13 aHC sites, 10 pHC sites). In this population with more normal sleep, for aHC, the mean SWR rate was 11.97/min in N2, and significantly different at 18.29/min in N3 (two-tailed paired t-test, p = 0.0089); for pHC, the mean SWR rate was 6.85/min in N2, and significantly different at 10.02/min in N3 (two-tailed paired t-test, p = 0.0166). Also, similarly to the overall results, under different NREM stages, the mean SWR rates in aHC were significantly different from those in pHC (two-tailed t-tests, p = 0.0306 for N2, p = 0.0191 for N3).

## Discussion

In this study, we applied rigorous selection criteria to arrive at a reliable population of human hippocampal sharpwave-ripples (HC-SWR) that differ from interictal spikes (IIS) and resemble rodent HC-SWR in key traits. We therefore expect our methods to aide future studies aiming to better understand HC-SWR’s contribution to human memory consolidation.

HC-SWR have previously been studied primarily in rodents (Buzsáki, 2015); human recordings are rare and limited to those obtained from patients with epilepsy. Generally, such studies have not clearly isolated HC-SWR from epileptiform activity, have failed to use rigorous criteria for identification of HC-SWR, have recorded from sites that were adjacent to the HC but seldom from the HC itself, and/or have not clearly defined the patient’s sleep/waking state (Table 1). Here, we selected patients whose hippocampal recordings appeared to be free of epileptic activity, and used rigorous selection criteria to clearly identify SWR. This allowed us to recognize SWR in a larger group of patients in HC with limited epileptiform activity; unlike sleep/waking state-independent pathological high-frequency activity found in humans (Staba et al., 2004) that spreads over NREM as well as REM/waking, the occurrence rate of our HC-SWR is highly concentrated in NREM. We then extensively characterized human HC-SWR so they could be compared to previous studies in rodents.

Human HC-SWR resemble rodent’s in many basic characteristics. In rodents, ripples are maximal in stratum pyramidale of CA1, and sharpwaves in stratum radiatum (Buzsáki, 2015). We observed a consistent topography from ripple to sharpwave maxima as the contacts progressed lateral to medially in humans.

Like most previous studies, we found a ripple center frequency between 80 and 90Hz, much lower than rodents (~140 Hz), but similar to monkeys (Skaggs et al., 2007; Logothetis et al., 2012; Ramirez-Villegas et al., 2015). Staba *et al*. (2004) found that bursts of power between 80-500Hz in human HC peaked at either 89 (ripple) or 263 (fast ripple). Both ripples and fast ripples occurred in the side contralateral to seizure onset, but the proportion of ripples was greater on that side; no requirement for an associated SW was imposed. We show that the lower center frequency characterizes ripples within sharpwaves, in HC without IIS. Ripples may comprise an exception to the general rule that frequencies of identifiable rhythms are constant across mammalian species (Buzsáki et al., 2013). The functional implications of the lower frequency in humans is unclear. In rodents, 80-90Hz oscillations are seen when the depolarization of CA1 pyramidal cells is less than during ripples (Sullivan et al., 2011). Alternatively, the lower frequency may reflect larger cells, lower packing density, and/or different channel time-constants in the human HC.

Previous reports of human ripple density range from 0.08/min (Staba et al., 2004) to 30/min, or even 115/min if patients with memory deficits are included (Jacobs et al., 2016) (Table 1). The average SWR density in this study ranged from ~7 to ~17/min, according to the stage of sleep or part of the hippocampus being recorded. This density was consistent across patients with no HC-IIS or seizures, versus those with some HC-IIS, and lies within the range of previous human (Table 1) and animal studies (Buzsáki, 2015).

Our finding that HC-SWR are preceded by suppression of broadband oscillations in the putative CA1 is consistent with models that emphasize a role for sustained inhibition in the genesis of SWR (Buzsáki, 2015). As in rodents (Klausberger et al., 2003), putative inhibitory interneurons in human hippocampus fire before SWR (Le Van Quyen et al., 2008). The quiet background may increase the consistency of CA1 neural response to CA3 input which triggers SWR.

The long, tubular human hippocampus shows substantial differences in anatomical connections and function along its length (Poppenk et al., 2013; Strange et al., 2014). While primate dorsal HC has been obliterated during fetal development, it is unclear if the remaining HC corresponds to the rodent ventral HC only, or if the HC has rotated such that human posterior HC corresponds to rodent dorsal.

In rats, SWR density depend on the interaction of threshold and location on the dorso-ventral HC axis: smaller ripples occurred at ~15/min in the ventral HC and 5 in dorsal, whereas larger ripples the converse (Patel et al., 2013). Our SWR results in humans were like the smaller ripples in rats: ~15/min for anterior and 8/min for posterior HC. Recently, Staresina *et al*. (2015), recording mainly in posterior HC, reported that human ripples were phase-coupled to HC spindles, rather than the sharpwaves typical in rodents. Alongside our results, this suggests that spindle-ripples rather than sharpwave-ripples may characterize posterior HC.

Human NREM differs from rodent’s, being nocturnal and quasi-continuous, and including distinct N2 and N3 stages. In each 90-min sleep cycle N2 precedes N3, but most N2 occurs later in the night, and N2/N3 may make complementary contributions to memory consolidation (Wei et al., 2018). Only one previous study reported different rates for ripples in N2 (1.6/min) versus N3 (1.0/min), but did not record from the HC, while explicitly including IIS with SWR (Clemens et al., 2007). In contrast, we found that human HC-SWR, excluding IIS, were lower density in N2, 9/min, than N3, 15/min. Traditionally, N2 is characterized by sleep spindles and K-complexes, and N3 by down-upstates (Silber et al., 2007), suggesting that SWR may be more strongly associated with down-upstates than spindles. However, spindles and down-upstates both occur in both stages (Mak-McCully et al., 2017), and both co-occur with SWR (Buzsáki, 2015). In any case, we show that human HC-SWR are present in both sleep stages.

In rodents, SWR are common during Quiet Waking (Roumis and Frank, 2015), a behavioral state characterized by grooming or immobility and large cortical LFP in comparison to active waking (Poulet and Petersen, 2008). In humans, Axmacher *et al*. (2008) reported ripples during quiet waking, at a density (5.3/min) much higher than N3 (0.35/min). In contrast, Clemens (Clemens et al., 2007) reported lower ripple density in waking (0.85/min) versus N2/N3 (1.3/min), but these recordings were extra-HC and included IIS. Finally, Brázdil *et al*. (2015), who recorded only during a task, reported high ripple density (19.4/min). We found that human HC-SWR are very rare during waking (1.9/min) compared to N2/N3 sleep (12/min). However, unlike the human studies cited above, we required that ripples be embedded in a sharpwave; removing that requirement led to greater density of ripples during waking than NREM (Fig. 3*A*-*D*). The occurrence of ripples without a sharpwave has also been noted in rodents and macaques (Buzsáki, 2015; Ramirez-Villegas et al., 2015). It is possible that ripples during waking (as opposed to SWR) contribute to consolidation in humans, and conversely, there is no direct evidence that SWR during NREM contribute to memory consolidation in humans. Axmacher *et al*. (2008) found that ripples in rhinal cortex, mainly recorded during quiet waking, were correlated with consolidation. More recent human intracranial EEG studies (Jiang et al., 2017; Zhang et al., 2018) found during NREM a significant co-occurrence of human HC-SWR with replay of widespread cortical spatiotemporal gamma/high gamma patterns from the preceding waking period.

Our study demonstrates that SWR occur in human hippocampi seemingly free from epileptiform activity, and that similar SWR occur in HC showing only effects of distant foci. The human HC-SWR identified by us resemble rodent’s in their waveforms, distribution across putative lamina in CA1, concentration in NREM sleep, and overall density, thus lending support to their being distinct from pathological phenomena. Overall, this study provides a framework for, and imposes powerful constraints on, models for the hippocampal contribution to memory consolidation in humans.

## Acknowledgements

This work was supported by the U.S. Office of Naval Research’s Multidisciplinary University Research Initiatives Program (N00014-16-1-2829), the National Institute of Biomedical Imaging and Bioengineering (R01 EB009282), and the National Institute of Mental Health (R01 MH111437). The authors would like to thank the following for their support: Qianqian Deng, Charles Dickey, Darlene Evardone, Zach Fitzgerald, Chris Gonzalez, Don Hagler, Milan Halgren, Erik Kaestner, Rachel Mak-McCully, Adam Niese, Burke Rosen, and T. G. Venti. The authors declare no conflict of interest.

